# TW68, Cryptochromes stabilizer, regulates fasting blood glucose level in *ob/ob* and fat-induced diabetic mice

**DOI:** 10.1101/2023.07.13.548861

**Authors:** Saliha Surme, Cagla Ergun, Seref Gul, Yasemin Kubra Akyel, Zeynep Melis Gul, Onur Ozcan, Ozgecan Savlug Ipek, Busra Aytul Akarlar, Nurhan Ozlu, Ali Cihan Taskin, Metin Turkay, Ahmet Ceyhan Gören, Ibrahim Baris, Nuri Ozturk, Mustafa Guzel, Cihan Aydin, Alper Okyar, Ibrahim Halil Kavakli

## Abstract

Cryptochromes (CRYs), transcriptional repressors of the circadian clock in mammals, inhibit cAMP production when glucagon activates G-protein coupled receptors. Therefore, molecules that modulate CRYs have the potential to regulate gluconeogenesis. In this study, we discovered a new molecule called TW68 that interacts with the primary pockets of mammalian CRY1/2, leading to reduced ubiquitination levels and increased stability. In cell-based circadian rhythm assays using U2OS:*Bmal1*-d*Luc* cells, TW68 extended the period length of the circadian rhythm. Additionally, TW68 decreased the transcriptional levels of two genes, *Phosphoenolpyruvate carboxykinase 1* (*PCK1*) and *Glucose-6-phosphatase* (*G6PC*), which play crucial roles in glucose biosynthesis during glucagon-induced gluconeogenesis in HepG2 cells. Oral administration of TW68 in mice showed good tolerance, a good pharmacokinetic profile, and remarkable bioavailability. Finally, when administered to fasting *ob/ob* and fat-induced diabetic animals, TW68 reduced blood glucose levels by enhancing CRY stabilization and subsequently decreasing the transcriptional levels of *Pck1* and *G6pc*. These findings collectively demonstrate the antidiabetic efficacy of TW68 *in vivo*, suggesting its therapeutic potential for controlling fasting glucose levels in the treatment of type 2 *diabetes mellitus*.

## Introduction

The circadian rhythm plays a crucial role in helping organisms adapt to the daily changes caused by the rotation of the Earth. In mammals, the circadian clock is synchronized by external cues such as light-dark cycles and food intake [1, 2]. Various physiological processes, including metabolism [3, 4], feeding behavior [5], sleep and wake cycles [6], locomotor activity [7] and hormonal activity [8] are regulated by the circadian rhythm. Previous studies using microarray and RNA-seq have demonstrated that more than 20% of genes in mammals are under the control of the circadian clock [9, 10]. Consequently, any disruption to the circadian clock mechanism can interfere with these physiological processes and contribute to the development of certain diseases, such as sleep, metabolic, and mood disorders [11, 12]. While light therapy, melatonin-like medications, and behavioral therapies are commonly used for treatment, there has been a recent focus on the discovery of small molecules that modulate the circadian rhythm for therapeutic purposes [13, 14].

The molecular mechanism of the mammalian circadian clock involves a transcriptional-translational feedback loop with positive and negative regulators. In the positive feedback loop, the transcription factors BMAL1 and CLOCK form a heterodimer that activates the expression of clock-controlled genes (CCGs) by binding to the E-box element within their promoters including *Per* and *Cry* [15, 16]. PER and CRY proteins accumulate in the cytosol and translocate into the nucleus together with casein kinase Iε (CKIε) [17–20]. The degradation of CRY and PER proteins relieves the repression of BMAL1/CLOCK, restarting the cycle anew [9]. Moreover, this mechanism is subject to fine-tuning by post-translational modifications such as ubiquitination, sumoylation, acetylation, and phosphorylation [21–24]. The stability of CRY1/2 proteins is primarily controlled by phosphorylation through AMPK, followed by polyubiquitination and degradation by the 26S proteasome [25]. E3 ubiquitin ligases, SKP1–CUL1–F-box protein complexes, SCF-FBXL3, and SCF-FBXL21, are involved in the degradation of CRYs. FBXL21-mediated degradation is much slower than that of its paralogue, FBXL3 [26, 27]. Consequently, FBXL21 increases the stability of CRY proteins by protecting them from faster degradation [28, 29]. The interaction between CRY1/2 and FBXL3/21 occurs through the FAD-binding pocket in the PHR domain [26, 30].

In recent years, there has been significant research focused on the discovery of small molecules that interact with circadian clock proteins to modulate their activity, with the aim of finding potential treatments for related disorders. These studies have led to the identification of several molecules that impact various aspects of the circadian rhythm [31–35]. For instance, a molecule called KL001, which was identified through high-throughput screening, has been found to increase the stability of CRY proteins and suppress gluconeogenesis in primary hepatocytes [36]. Subsequent studies involving derivatization based on KL001 have led to the identification of certain isoform-specific CRY-targeting molecules [37, 38]. However, there are no available reports about their efficacy in animals.

In this study, our aim was to identify a small molecule that binds to CRY proteins by targeting their critical primary pocket involved in the interaction with FBXL3. We employed a structure-based approach, starting with virtual screening of a library of approximately 2 million small molecules. Through subsequent biochemical and molecular evaluations, we narrowed down the selection of molecules. *In vitro* studies revealed a CRY-binding small molecule called TW68, which was found to stabilize both CRY1 and CRY2. Using cell-based circadian rhythm analysis, we observed that TW68 increased the period length in U2OS:*Bmal1*-d*Luc* cells. Furthermore, treatment of HepG2 cell lines with TW68 resulted in a reduction in glucose levels due to decreased mRNA levels of the *PCK1* (*Phosphoenolpyruvate Carboxykinase 1*) and *G6PC* (*Glucose-6-phosphatase*) genes, which play crucial roles in gluconeogenesis. After conducting toxicity studies and determining the pharmacokinetic profiles of TW68 in mice, we tested its effect on blood glucose levels in diabetic animals, including genetically obtained *ob/ob* mice and fat-diet-induced diabetic animals. The results demonstrated that TW68 decreased fasting blood glucose levels in both animal models. Finally, we showed that TW68 increases the stability of CRY1/2 and reduces the transcription of the *Pck1* and *G6pc* in mice liver.

Overall, our findings indicate that TW68 possesses the ability to increase the stability of CRY1/2, reduce the transcription of *Pck1* and *G6pc*, and lower fasting blood glucose levels *in vivo*.

## Results

### In-silico Screening and Cell Viability Assessment

CRY proteins are characterized by three functionally important domains: the extended C-terminal region, the primary pocket, and the secondary pocket [39, 40]. The primary pocket is located in the α-helical domain, while the secondary pocket is found in the α/β domain. The C-terminal region, although disordered, plays a crucial role in interacting with various proteins, including CBS [39, 41]. The primary pocket of CRYs is particularly significant as it is involved in the interaction with FBXL3, an E3 ligase that regulates their stability [42]. Thus, we focused on this site for virtual screening to identify novel molecules capable of modulating CRY stability, as described in [43]. For the virtual screening, we utilized commercially available small molecule libraries (Ambinter (Orleans-France)) that had unknown functions. These compounds were docked into the primary pocket of CRY1 using computational techniques. We calculated the binding energies of the compounds, and based on their scores, we selected the top compounds with the lowest binding energies to CRY1 (ranged from −9 to −13.5 kcal/mol). These compounds were deemed the most promising candidates for further experimental studies. Molecules to be tested were commercially purchased from Ambinter (Orleans-France). Initially, we performed a toxicity assessment of selected molecules using U2OS cells. The cells were treated with each molecule at varying concentrations, and the viability of the cells was measured using the MTT assay. Non-toxic dosages of molecules were used for further characterization.

### The Effect of Small Molecules on CRY1/2 Half-life

We utilized various *in vitro* tools based on the luciferase system. To eliminate molecules that affect luciferase activity or stability, we assessed the impact of these molecules on luciferase activity. HEK 293T cells were transfected with plasmids containing the d*Luc* gene and treated with small molecules at their non-toxic concentrations. Molecules affecting the luciferase half-life were identified and excluded from the further investigation. Since our goal was to identify molecules that regulate the half-life of the CRYs, we subsequently evaluated the effect of remaining small molecules on the half-life of both CRY1 and CRY2 by measuring the decay rates of CRY1:LUC and CRY2:LUC (where the C-terminal of CRYs was fused with luciferase). To this end, HEK293T cells were transfected with pcDNA-*Cry1-Luc* or pcDNA-*Cry2-Luc* plasmids. Cells were, then, treated with cycloheximide (CHX) in the presence of the small molecules. Small molecule named as TW68 enhanced the half-life of CRY1 and CRY2, and affected the circadian rhythm of U2OS *Bmal1*:d*Luc* cells in a dose-dependent manner (**Fig. 1C**). TW68 was then subjected to further characterization.

**Figure 1:**
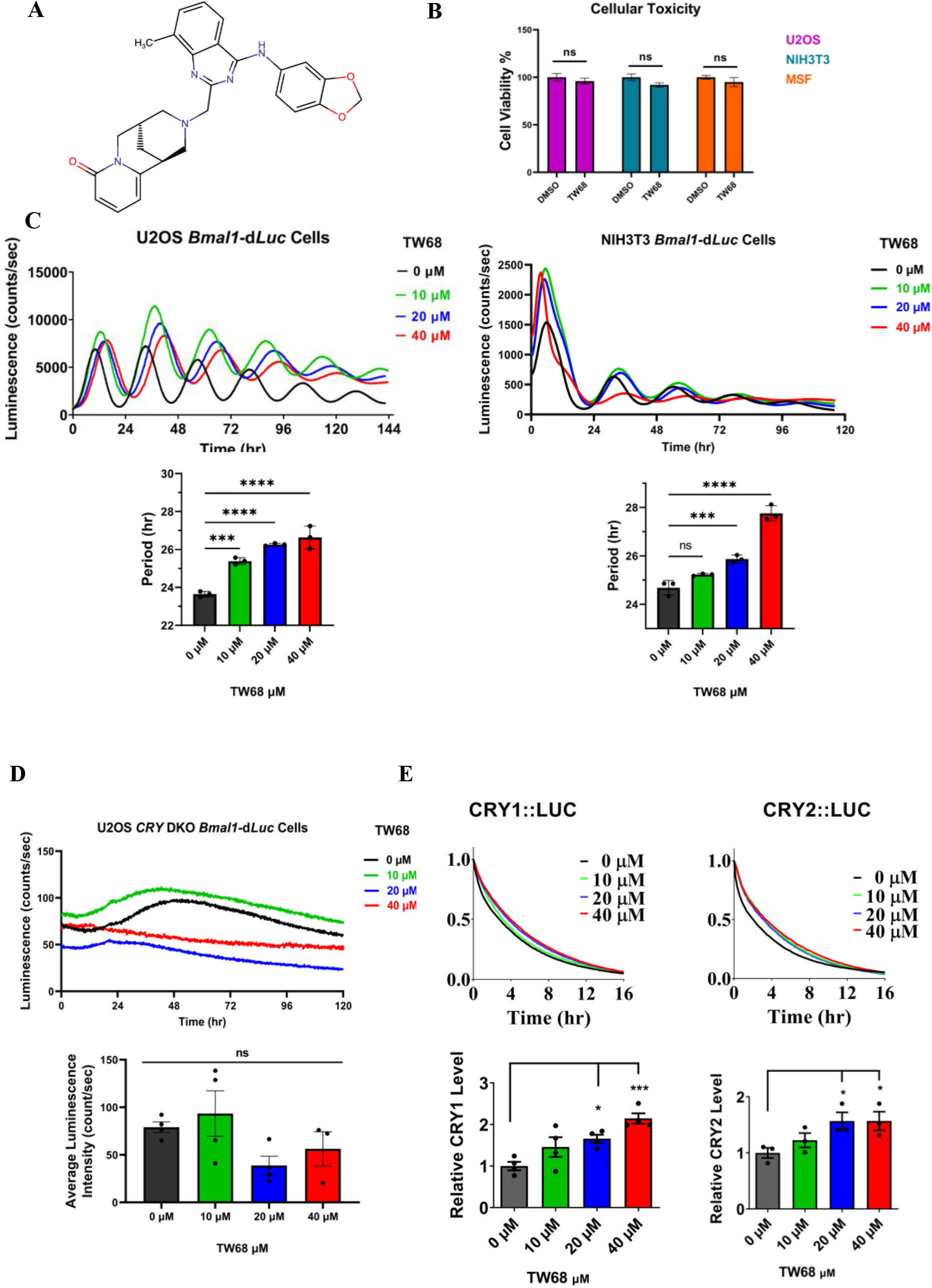
The effect of TW68 on cell toxicity and circadian rhythm. A) chemical structure of TW68. B) The toxicity analysis of TW68 on U2OS, NIH 3T3 and primary mouse fibroblast (MSF) cell lines. Experiments were performed using MTT assay at 20 µM TW68. Data represent the mean ± SEM, n = 3. DMSO control versus TW68 treated cells by student-test (ns=not significant). C) The effect of TW68 on circadian rhythm using U2OS and NIH3T3 stably expressing destabilized luciferase under the control of *Bmal1* promotor. (Data represent the mean ± SEM, n = 3DMSO control versus TW68 treated cells by one-way ANOVA with Dunnett’s multiple comparisons tests **p<0.01, ***p<0.001, ****p<0.0001). D) The effect of TW68 on the *CRY1/CRY2* double knockout (DKO) U2OS cell line (Data represent the mean ± SEM, n = 3. DMSO control versus TW68 treated cells by one-way ANOVA with Dunnett’s multiple comparisons test ns=nonsignificant). E) TW68 enhances the CRY1 and CRY2 half-life in a dose-dependent manner Normalized half-life is shown with ± SEM, n = 3. DMSO control versus TW68 treated cells by one-way ANOVA with Dunnet’s multiple comparison test (*p<0.05, **p<0.01).

### Characterization of the TW68

We evaluated the cytotoxicity of TW68 (**Fig. 1A**) on U2OS, NIH 3T3 and mouse primary skin fibroblast cells (MSF) at a concentration of 20 µM using the MTT assay. Our findings demonstrated that TW68 did not exhibit any toxic effects on commonly utilized cell lines involved in circadian rhythm research, as well as the primary MSF cell line (**Fig. 1B**). Subsequently, we investigated the impact of TW68 on circadian rhythm using U2OS *Bmal1*:d*Luc* cells and NIH 3T3 *Bmal1:*d*Luc* cells. The cell lines stably express destabilized luciferase under the control of *Bmal1* promotor and are used in various studies for the screening of the molecules [32–34, 36, 38]. The analyses of the circadian rhythm revealed that TW68 lengthened the period in both cell lines (**Fig. 1C**). To eliminate the possibility that TW68 might affect other proteins and, in turn, circadian rhythmicity, we tested its effect on the *CRY1* and *CRY2* double knockout U2OS *Bmal1*:d*Luc* cell line generated by Gul et al. [33]. TW68 did not affect luminance in a dose-dependent manner in the absence of CRYs, implying that the effect of TW68 was via CRYs (**Fig. 1D**). Based on the dose-response of the CRY1-LUC half-life, the EC_50_ value of TW68 was estimated to be ∼10 µM.

We next determined the effect of the TW68 on the half-life of CRYs by determining the decay rates of CRY1::LUC and CRY2::LUC as described in [36]. For this purpose, HEK 293T cells were transfected with pcDNA-*Cry-Luc* plasmids and treated with TW68 in a dose-dependent manner. The next day, cells were treated with CHX, and luminescence decay was recorded. Analysis showed that TW68 increased the half-life of CRY1 and CRY2 in a dose-dependent manner (**Fig. 1E**).

The docking poses of TW68 indicate that it binds to the primary pockets of CRY1 and CRY2 (**Fig. 2A**) by interacting with specific amino acid residues: Arg358 (Arg376 in CRY2), Ser396 (Ser 414 in CRY2), His359 (His377 in CRY2), and Trp399 (Trp417 in CRY2) of CRY1 (**Fig. 2B**). The calculated binding energies between TW68 and CRY1 and CRY2 were −9.7 kcal/mol and −10.4 kcal/mol, respectively (**Fig. 2C**). To confirm the binding of TW68 binds this region, biotinylated TW68 (bTW68) (**Fig. 3A**) was synthesized. Initially, the activity of the bTW68 was shown on circadian rhythm of U2OS *Bmal1*:d*Luc*. Then, to show the physical interaction between CRYs and bTW68, pcDNA-*Cry1-His-Myc* and pcDNA-*Cry2-His-Myc* were transfected into HEK293T cells. After the harvesting of the cells and the preparation of the cell lysates, bTW68 was mixed with them for the pull-down of CRY1-HIS-MYC and CRY2-HIS-MYC in the presence and absence of the competitor (free TW68). As shown in **Fig. 3B**, both CRYs were precipitated with bTW68, but the level of CRYs was significantly reduced in the presence of the competitor. This suggests that TW68 physically interacts with both CRYs. Crystal structural studies with KL001 were shown to bind the primary pocket of CRY1 [44]. We reasoned that because TW68 was computationally shown to bind to the primary pocket of CRYs, KL001 and TW68 should compete for the same binding site. Therefore, we performed a pull-down assay for bTW68 with and without free KL001. The binding of bTW68 was greatly reduced in the presence of KL001 (**Fig. 3C**). All these results suggested that TW68 physically interacts with both CRYs through the primary pocket. Next, to investigate the off-target effect of TW68 we used bTW68 for the pull-down study using protein extract prepared from the HEK293T cell line in the presence and absence of the competitor (TW68). After pulldown, resin was used for the LC-MS/MS analysis as described by in [36]. The results were given in **Table 1**. The bTW68 binds to CRY1, and its interaction reduced when competitors were present. The other three identified proteins in **Table 1** were determined to be sticky proteins from the Crapome Database.

**Figure 2:**
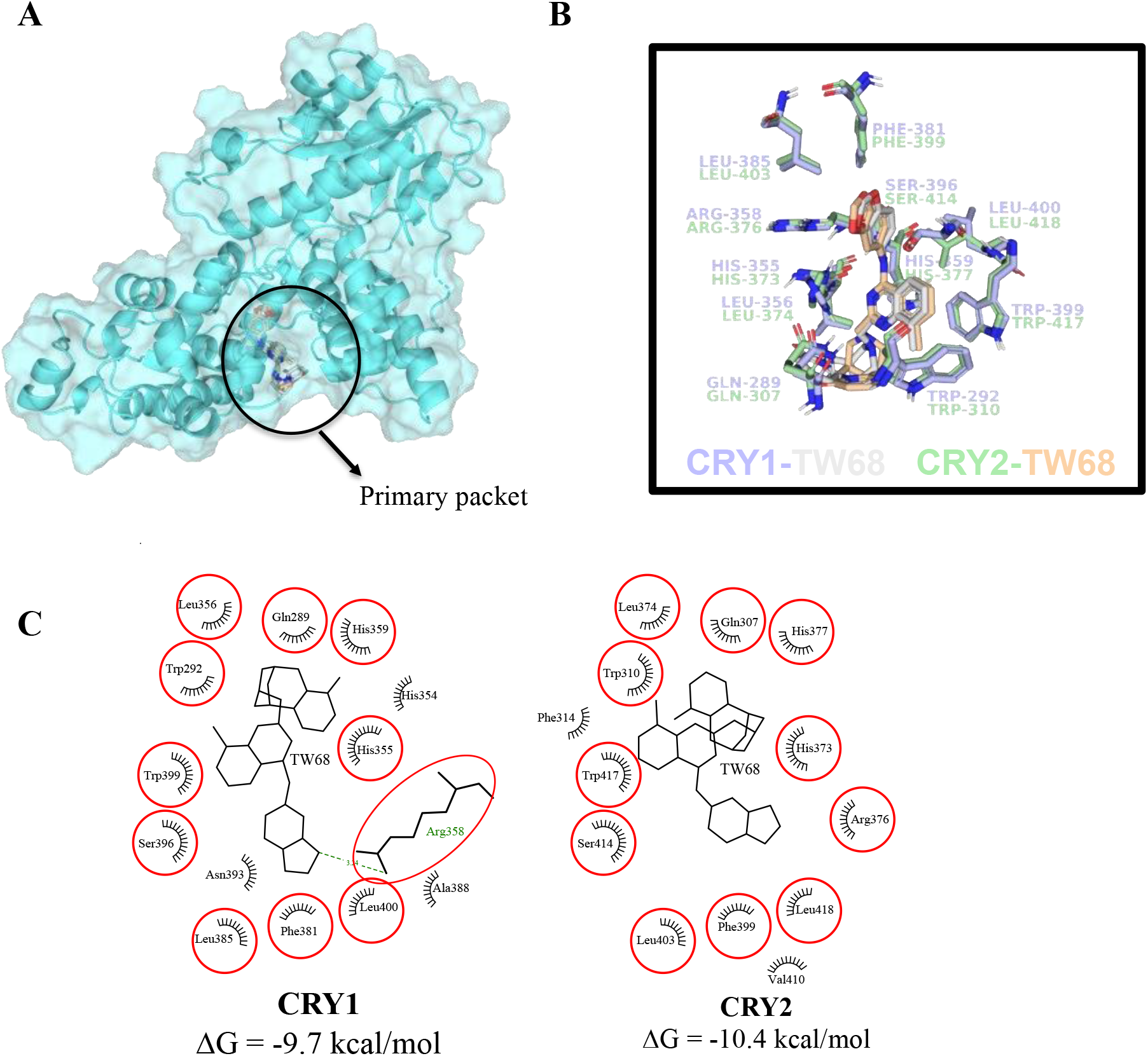
Comparative analysis of interactions that TW68 forms with CRY1 and CRY2. (A) TW68 docked into primary pocket of CRY1 PHR (PDBID:6KX7) and CRY2 PHR domains (PDBID:6KX8) using Autodock Vina. TW68 docked halo models were superimposed and TW68 from CRY1 and CRY2 were colored grey and yellow, respectively. CRY1 PHR domain (PDBID:6KX7) used for representation. (B) Common amino acid residues that are in close contact with TW68 in both CRY1 and CRY2 are represented as purple and green sticks, respectively. (C) 2D interaction diagram of TW68 with CRY1 (left) and CRY2 (right) were plotted in LigPlot+. Equivalent residues that are in interaction with TW68 in both CRY1 and CRY2 are encircled in red. ΔG values are respective scores from molecular docking in Autodock Vina.

**Figure 3:**
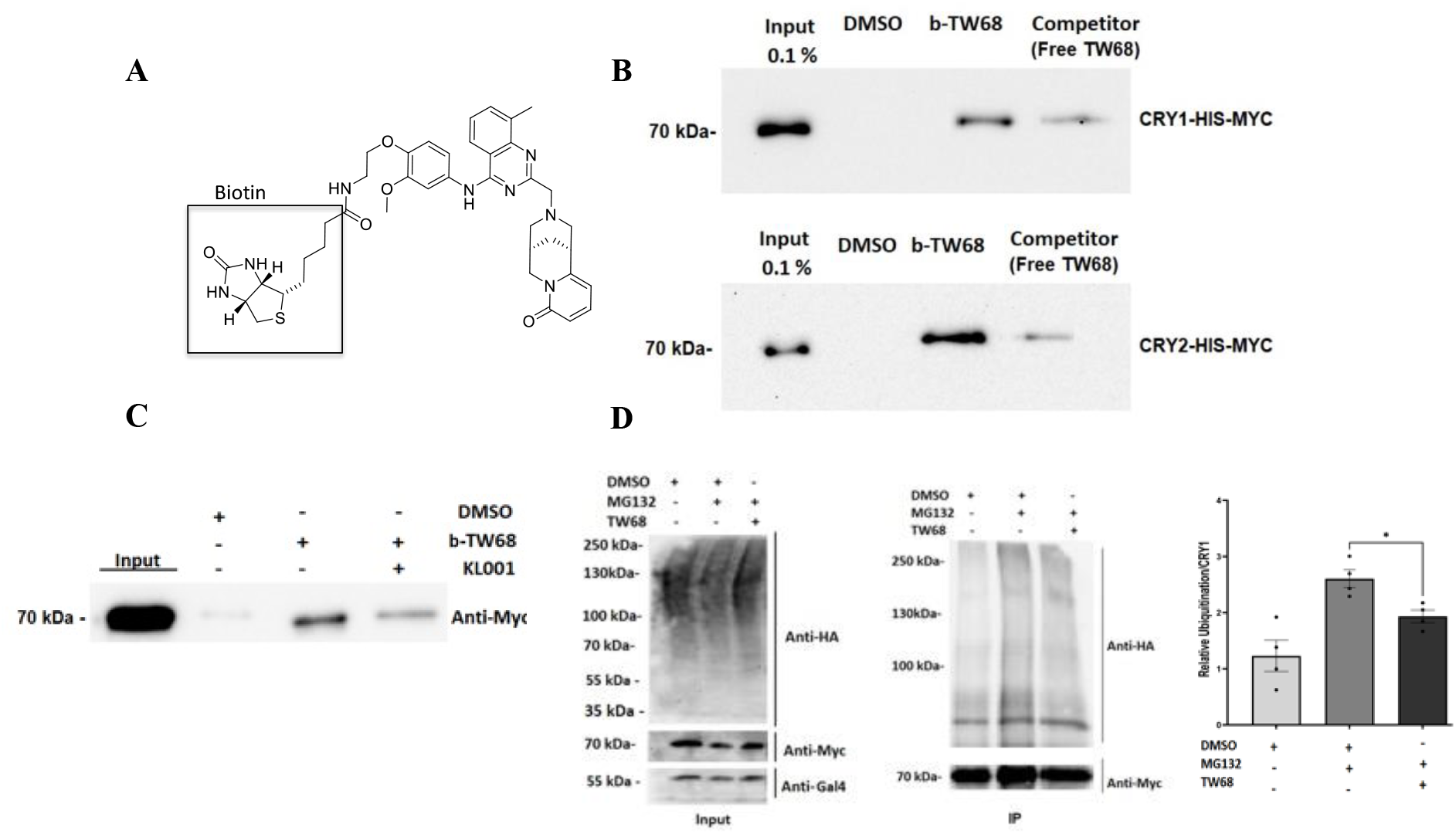
The physical interaction between TW68 and CRYs. A) The chemical structures of biotinylated-TW68 (bTW68). B) bTW68 was used in to pull down CRY1 and CRY2 from HEK 293T cell extract transfected with pcDNA-*Cry1-His-Myc* and pcDNA-*Cry2-His-Myc*. While bTW68 binds to both CRYs, its interaction is reduced in the presence of TW68 (competitor). C) KL001 and TW68 compete for the same binding site. The binding of bTW68 was highly reduced in the presence of KL001. D) The representative figure of the ubiquitination of CRY1 in the presence and absence of TW68. (Data represent the mean ± SEM, n =4; *p<0.05 by DMSO control with unpaired t-test with two-tailed).

**Table 1:** Identification of TW68 binding proteins using LC-MS/MS

| Gene Name | Uniprot ID | Gene Symbol | DMSO |  |  | TW68-Biotinylated |  |  | Competitor |  |  | CRAPOME |  |  |
| --- | --- | --- | --- | --- | --- | --- | --- | --- | --- | --- | --- | --- | --- | --- |
|  |  |  | Rep1 | Rep2 | Rep3 | Rep1 | Rep2 | Rep3 | Rep1 | Rep2 | Rep3 | Num of Expt. (found/total) | Ave SC | Max SC |
| Cryptochrome-1 OS=Homo sapiens GN=CRY1 PE=1 SV=1 | Q16526 | CRY1 | 0 | 0 | 0 | 12 | 11 | 8 | 0 | 0 | 0 | 7 / 716 | 0 |  |
| Exportin-1 OS=Homo sapiens GN=XPO1 PE=1 SV=1 | O14980 | XPO1 | 0 | 0 | 0 | 8 | 10 | 6 | 0 | 0 | 2 | 237 / 716 | 3.8 | 52 |
| Tubulin beta-4A chain OS=Homo sapiens GN=TUBB4A PE=1 SV=2 | P04350 | TUBB4A | 0 | 0 | 0 | 16 | 16 | 10 | 0 | 2 | 0 | 668 / 716 | 30.7 | 255 |
| Tubulin alpha-1A chain OS=Homo sapiens GN=TUBA1A PE=1 SV=1 | Q71U36 | TUBA1A | 0 | 0 | 0 | 19 | 12 | 12 | 7 | 0 | 0 | 694 / 716 | 41.7 | 261 |

Since TW68 binds to the primary pockets of CRYs, which is an important region for the binding of FBXL3 (E3 ligase) and degradation of CRYs by causing their ubiquitination and subsequent proteasomal degradation [45–47], we aimed to investigate the effect of TW68 on the ubiquitination levels of CRYs. To test this, HEK293T cells were transfected with pcDNA-*Cry1*-His-Myc, *Fbxl3*-GAL4, and pUb-HA plasmids and then treated with TW68 or DMSO. Then, MG132 was added to each cell lysate to inhibit proteasomal degradation. Proteins were subjected to a pull-down assay using anti-MYC resin, and the ubiquitination levels of CRY1 were measured by Western blot. The results revealed that TW68 treatment led to a reduction in the ubiquitination of CRY1 compared to the DMSO control (**Fig. 3D**). It has been reported that the last five amino acid residues of the C-terminal tail of FBXL3 bind to the primary pocket of CRY1 and are crucial for CRY ubiquitination [48]. Therefore, TW68 likely disrupts the CRY-FBXL3 interaction, resulting in reduced ubiquitination and stabilization of CRYs.

All these results showed that TW68 interacts with the primary pocket in the PHR of CRYs and specifically stabilizes them by reducing their ubiquitination.

#### The effect of TW68 on the unsynchronized U2OS Cells

To evaluate the biochemical effect of TW68, unsynchronized U2OS cells were treated with the molecule in a dose-dependent manner. Cell lysates were prepared, and the samples were subsequently analyzed using SDS-PAGE followed by Western blotting with anti-CRY1, anti-CRY2, and anti-PER2 antibodies. The analysis revealed a significant dose-dependent increase in the levels of CRY1, CRY2, and PER2 compared to cells treated with DMSO as a control (**Fig. 4A**). This result suggested that TW68 stabilized both CRYs, which agrees with the CRY:LUC degradation assays. Next, we investigated the effect of TW68 on the transcription of clock output genes in these cells. Levels of clock gene transcripts were determined by qPCR. The tested gene mRNA levels were normalized with respect to *RPLP0* gene expression because its expression level is independent of the circadian rhythm. In agreement with the stabilization of CRYs, TW68 treatment resulted in a significant decrease of *CRY1*, *CRY2*, *PER2*, *DBP* and *REV-ERBα* expression in a dose-dependent manner (**Fig. 4B**). Furthermore, there was no significant effect of TW68 treatment on *BMAL1, RORα* and *CLOCK* expression (**Fig. 4B**). These results are consistent with the effects of CRY1/2 stabilization since the CRYs repress the expression of E-box regulated *PER2* and *DBP*.

**Figure 4:**
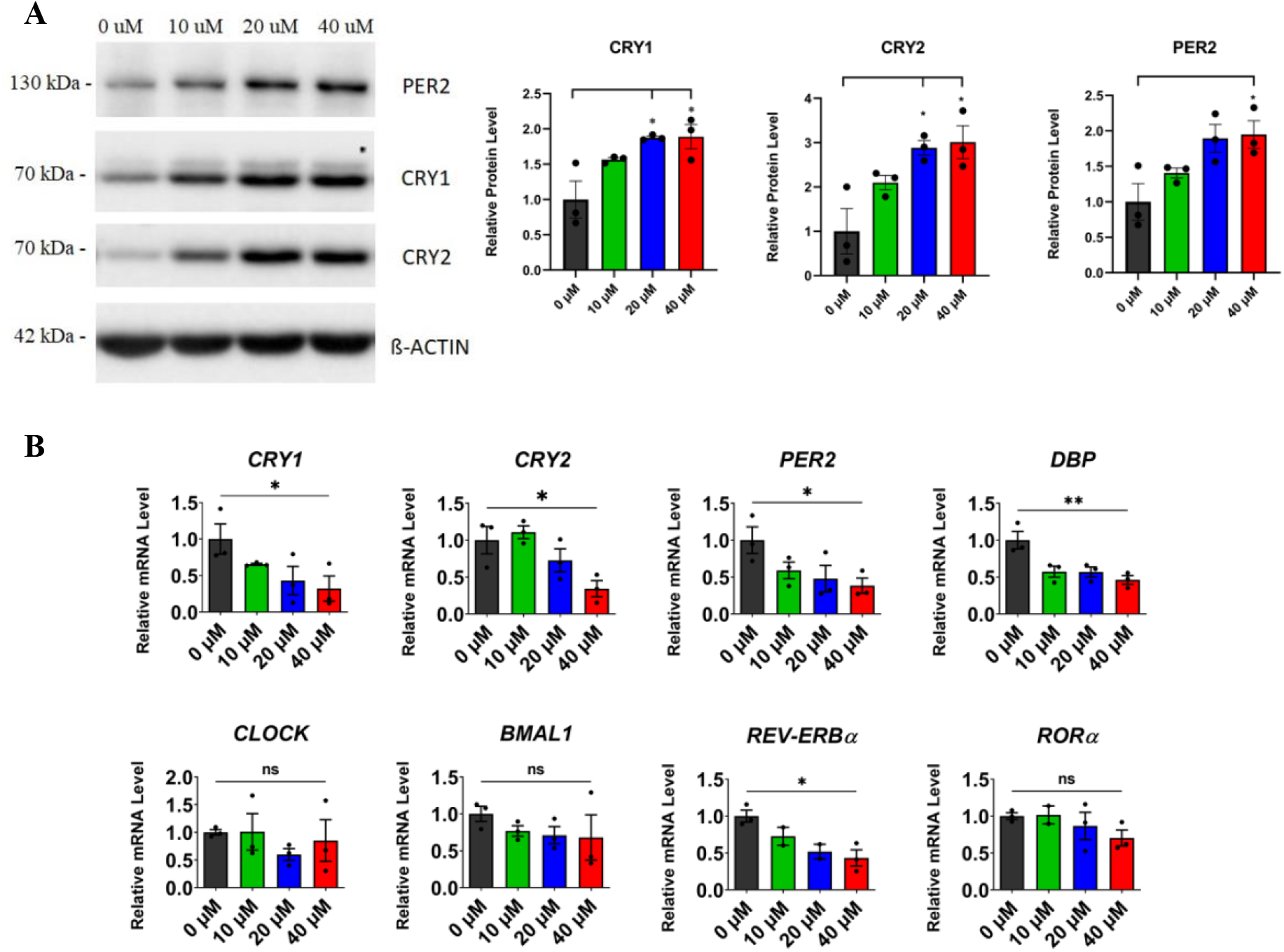
The dose-dependent effect of TW68 on core clock proteins and genes in unsynchronized U2OS cells. Unsynchronized U2OS cells treated with TW68 in dose-dependent manner. Cells are collected and analyzed for A) core clock protein level by Western blot (data represent the mean ± SEM, n =3; *p<0.05 by DMSO control one-way ANOVA with Dunnet’s multiple comparison test) B) the transcriptional level of core clock genes (data represent the mean ± SEM, n =3; *p<0.05 and **p<0.01 by DMSO control with one-way ANOVA with Dunnet’s multiple comparison test).

### TW68 Regulates Glucagon-induced Gluconeogenesis in HepG2 Cells

A Previous study has demonstrated that the stabilization of CRYs by a molecule with stabilizing properties can lead to a reduction in glucagon-mediated gluconeogenesis through its interaction with the Gsα subunit of heterotrimeric G-protein[49]. To investigate the effects of TW68 on glucagon-induced gluconeogenesis, HepG2 cells were treated with TW68, and the transcriptional levels of key genes involved in gluconeogenesis, phosphoenolpyruvate carboxykinase 1 (*PCK1*) and glucose-6-phosphatase (*G6PC*), were measured by qPCR. As shown in **Fig. 5A** the expression levels of *PCK1* and *G6PC* were significantly reduced upon exposure to TW68. Furthermore, the levels of glucose secreted by the HepG2 cells were measured, and the results indicated a dose-dependent reduction in glucose levels upon treatment with glucagon (**Fig. 5B**). Collectively, these findings suggest that TW68 stabilizes CRYs, subsequently leading to a decrease in glucagon-mediated gluconeogenesis.

**Figure 5.**
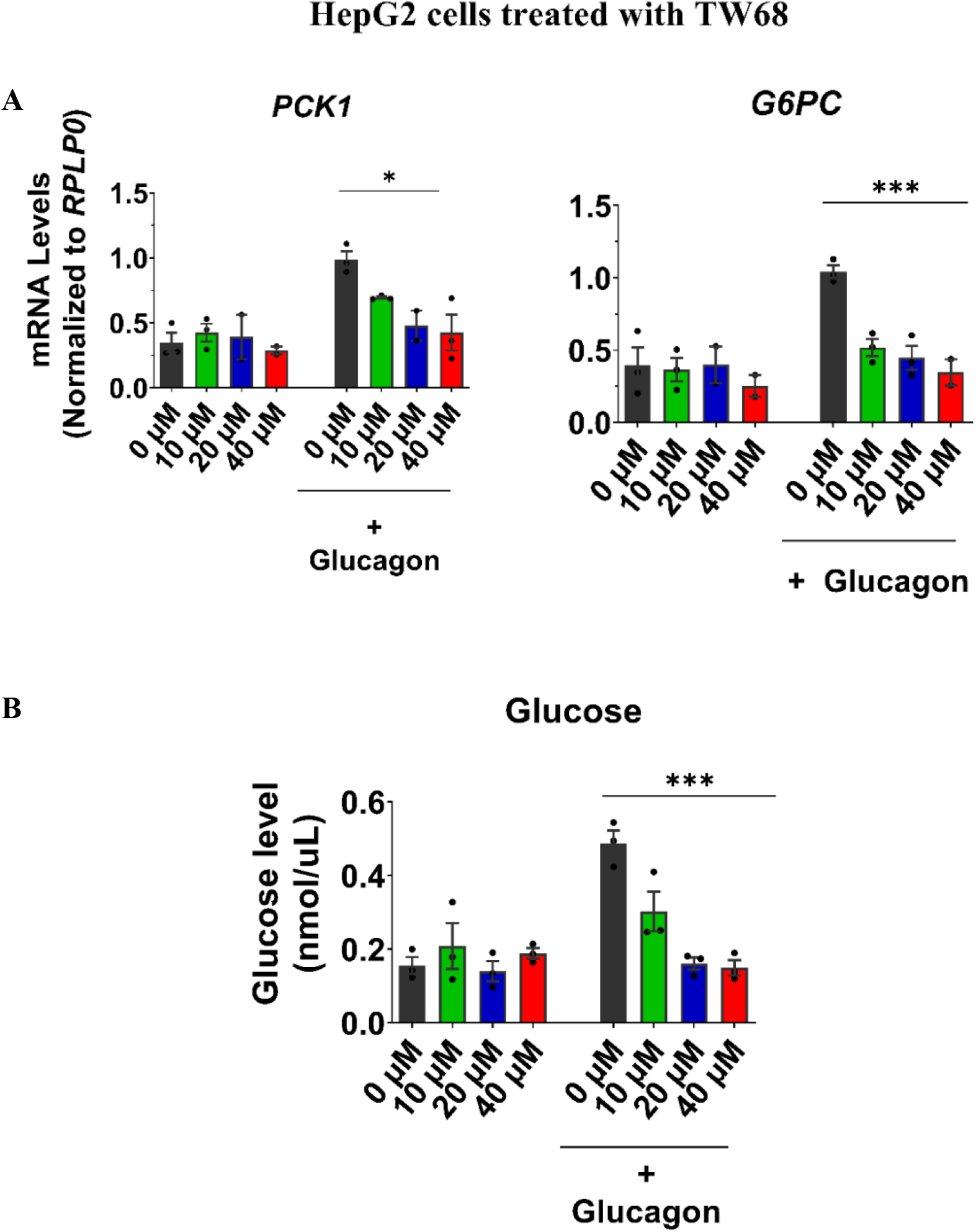
Effect of TW68 on gluconeogenesis in HepG2 cells. A) RT-qPCR analysis of gluconeogenic *PCK1* and *G6PC* genes when cells were treated with TW68 and induced or uninduced with glucagon**. B)** Glucose production amount of cells treated with TW68 and then induced or uninduced with glucagon. Statistical analysis was performed using one-way ANOVA (*p<0.05, ***p<0.001).

### Toxicity Studies of TW68 in Mice

To assess the single-dose toxicity (SDT) of TW68, C57BL/6J mice of both sexes were intraperitoneally (i.p.) administered with TW68 at doses of 50 mg/kg and 300 mg/kg. Following parameters were monitored to evaluate its toxicity: food and water consumption, behavior assessment, body weight changes, clinical signs, mortality, body temperature, and gross findings at terminal necropsy. Mice treated with 300 mg/kg of TW68 exhibited severe hypothermia, piloerection, a hunched posture, reduced locomotor activity, hyporeflexia, and dyspnea. These clinical signs indicated significant toxicity, leading us to consider the 300 mg/kg dose as lethal, and the animals were euthanized accordingly. Conversely, mice treated with 50 mg/kg of TW68 did not display any abnormal clinical signs compared to the control group. The body weight changes and body temperatures of these mice remained comparable to the control mice, with body temperatures ranging from 36.0-37.0 °C, indicating that a single dose of 50 mg/kg TW68 was well-tolerated (**Fig. 6A**). Overall, these findings suggest that the administration of a single dose of 50 mg/kg TW68 in mice was safe and did not induce significant adverse effects, while a higher dose of 300 mg/kg exhibited severe toxicity leading to the termination of the animals.

**Figure 6:**
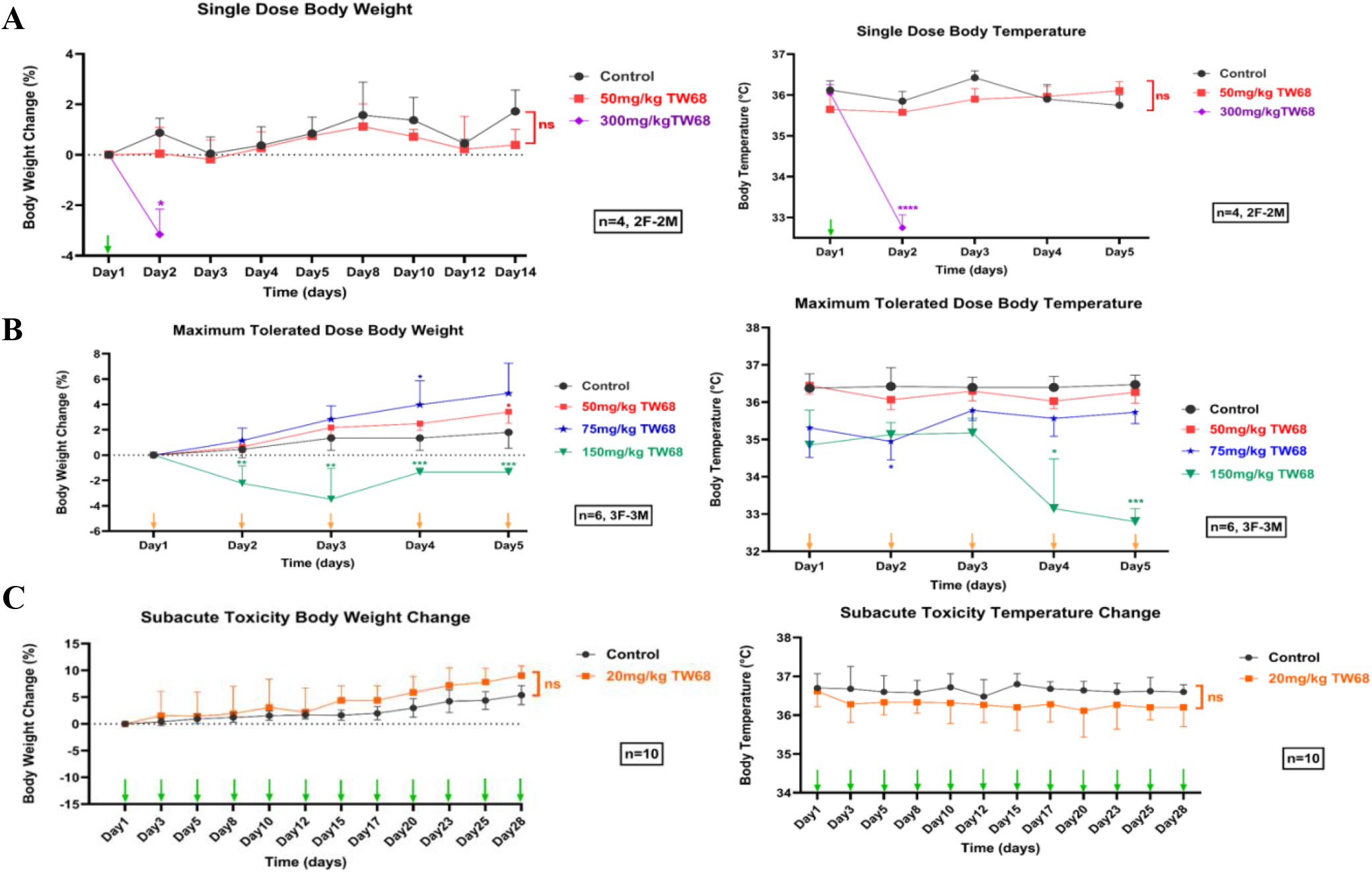
The toxicity of TW68 in mice A) Single Dose Toxicity of TW68 in mice. Body weight changes (%) in C57BL/6J mice treated with single intraperitoneal doses of 50 mg/kg, 300 mg/kg TW68 or Control vehicle during 14-day observation period. Body weight changes (%) were expressed as mean ± SEM. “↓” indicates the treatment days of TW68. *p<0.05, **p<0.01, ***p<0.001, **** p<0.0001 (Two-way ANOVA with Bonferroni post hoc test). Body temperatures in C57BL/6J mice treated with single intraperitoneal doses of 50 mg/kg, 300 mg/kg TW68 or Control vehicle during 14-day observation period. Body temperatures (°C) were expressed as mean ± SEM. “↓” indicates the treatment days of TW68. *p<0.05, **p<0.01, ***p<0.001, **** p<0.0001 (Two-way ANOVA with Bonferroni post hoc test). B) Maximum tolerated dose of TW68 in mice. Body weight changes (%) in C57BL/6J mice treated with intraperitoneal doses of 50 mg/kg, 75 mg/kg, 150 mg/kg TW68 or Control vehicle for 5 days. Body weight changes (%) were expressed as mean ± SEM. “↓” indicates the treatment days of TW68. *p<0.05, **p<0.01, ***p<0.001, **** p<0.0001 (Two-way ANOVA with Bonferroni post hoc test). Body temperatures in C57BL/6J mice treated with intraperitoneal doses of 50 mg/kg, 75 mg/kg, 150 mg/kg TW68 or Control vehicle for 5 days. Body temperatures (°C) were expressed as mean ± SEM. “↓” indicates the treatment days of TW68. *p<0.05, **p<0.01, ***p<0.001, **** p<0.0001 (Two-way ANOVA with Bonferroni post hoc test). C) Subacute Toxicity of TW68 in mice. Body weight changes (%) in C57BL/6J mice treated with intraperitoneal dose of 20 mg/kg TW68 or Control vehicle for 28 days. Body weight changes (%) were expressed as mean ± SEM. “**↓**” indicates the treatment days of TW68. *p<0.05, ***p<0.001, control vs 60 mg/kg M47 (Two-way ANOVA with Bonferroni post hoc test). Body temperatures in C57BL/6J mice treated with intraperitoneal dose of 20 mg/kg TW68 or Control vehicle for 28 days. Body temperatures (°C) were expressed as mean ± SEM. “**↓**” indicates the treatment days of TW68. *p<0.05, **p<0.01, ***p<0.001 (Two-way ANOVA with Bonferroni post hoc test).

The maximum tolerated dose (MTD) of TW68 over a 5-day period was determined by administering doses of 50, 75, or 150 mg/kg i.p. injections. Animals treated with 50 and 75 mg/kg of TW68 showed no observed adverse effects. However, when animals were treated with 150 mg/kg of TW68, three animals died by the end of the third day, resulting in a mortality rate of 66%. Changes in body weight were observed in animals treated with all three doses (**Fig. 6B**). Particularly, animals treated with 150 mg/kg of TW68 experienced weight loss (1-3%) throughout the duration of treatment. Conversely, animals treated with 50 and 75 mg/kg of TW68 showed weight gain on different days (**Fig. 6B**). Additionally, daily measurement of body temperature revealed significant hypothermia (32.4-34.8 °C) in animals treated with 150 mg/kg of TW68 compared to the control group (**Fig. 6B**). The dose of 75 mg/kg of TW68 caused mild hypothermia (35.0-35.8 °C). Animals treated with 50 mg/kg of TW68 showed no significant changes in body temperature compared to the control group. These results suggest that repeated doses of 50 mg/kg or below were well-tolerated by mice. Therefore, a subacute toxicity test was conducted using a dose of 20 mg/kg of TW68, administered daily for 28 days. No observable clinical symptoms were noted in these animals, and both body weight and temperature changes were comparable to the control animals treated with the vehicle throughout the 28-day period (**Fig. 6C**).

### Pharmacokinetic Profile of TW68 in Plasma and Tissues

To evaluate the pharmacokinetic properties of TW68, animals were intraperitoneally (i.p.) administered with a dose of 75 mg/kg. Following euthanasia, plasma and tissues (liver, kidney, and ileum) were collected at various time points to measure the concentration of TW68 over time. The mean concentration-time curves of TW68 in plasma and tissues of female C57BL/6J mice are presented in **Fig. 7A**, and the corresponding pharmacokinetic parameters are provided in **Table 2 and 3**. TW68 exhibited peak plasma and tissue concentrations at 0.5 hours after administration. The plasma half-life of TW68 was calculated as 1 hour. In addition, we also investigated the oral administration of TW68 to assess its absorption and bioavailability. Following oral administration, TW68 reached maximum levels in plasma and tissues at 0.5 hours (**Table 2 and 3**) and exhibited a plasma elimination half-life of 3.4 hours. Notably, TW68 demonstrated higher bioavailability compared to parenteral administration. Moreover, the tissue levels of orally administered TW68 were also higher compared to i.p. administration (**Fig. 7B**). Based on these findings, we decided to proceed with oral administration of TW68 to evaluate its efficacy in mice.

**Figure 7:**
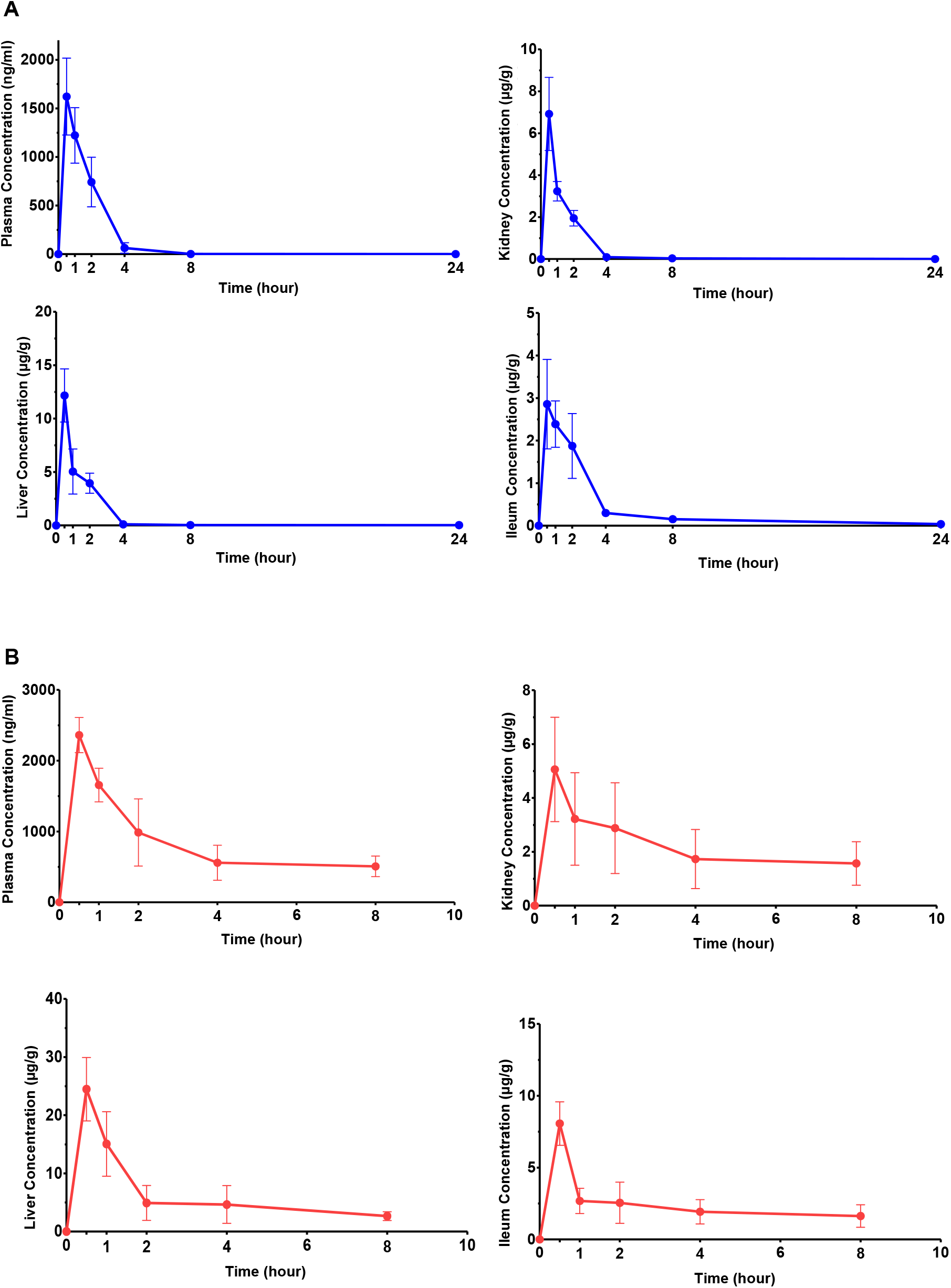
Pharmacokinetics of TW68 in mice A) Mean plasma and tissue concentration-time curve of TW68 administered at 75 mg/kg intraperitoneally to C57BL/6J mice. Data were expressed as mean ± SEM (n=4 per time point). B) Mean plasma and tissue concentration-time curve of TW68 administered at 75 mg/kg orally to C57BL/6J mice. Data were expressed as mean ± SEM (n=4 per time point).

**Table 2:** Plasma pharmacokinetic parameters of intraperitoneal (i.p.) and oral (p.o.) administered TW68 (75 mg/kg, single dose) Data were expressed as mean ± SEM (n=4 per time point).

| Parameters | Plasma (i.p.) | Parameters | Plasma (p.o.) |
| --- | --- | --- | --- |
| **C<sub>max</sub> (ng/ml)** | 1650,22 ± 368,35 | **C<sub>max</sub> (ng/ml)** | 2365,13 ± 251,50 |
| **t<sub>max</sub> (h)** | 0,5 | **t<sub>max</sub> (h)** | 0,5 |
| **AUC<sub>0-24</sub> (ng.h/ml)** | 3046,00 ± 915,70 | **AUC<sub>0-24</sub> (ng.h/ml)** | 5464,50 ± 1398,20 |
| **AUC<sub>0-∞</sub> (ng.h/ml)** | 3046,00 ± 915,70 | **AUC<sub>0-∞</sub> (ng.h/ml)** | 8503,94 ± 2601,52 |
| **t<sub>1/2</sub> (h)** | 1,02 ± 0,03 | **t<sub>1/2</sub> (h)** | 3,41 ± 0,92 |
| **CL (ml/h)** | 708,58 ± 247,47 | **CL/F (ml/h)** | 306,12 ± 148,37 |
| **Vd (ml)** | 1072,63 ± 404,63 | **Vd/F (ml)** | 1066,86 ± 220,49 |

**Table 3:** Tissue pharmacokinetic parameters of intraperitoneal (i.p.) and oral (p.o.) administered TW68 (75 mg/kg, single dose) Data were expressed as mean ± SEM (n=4 per time point).

| Parameters | Liver (i.p.) | Kidney (i.p.) | Ileum (i.p.) | Liver (p.o.) | Kidney (p.o.) | Ileum (p.o.) |
| --- | --- | --- | --- | --- | --- | --- |
| **C<sub>max</sub> (µg/ml)** | 12,16 ± 2,49 | 6,92 ± 1,75 | 2,98 ± 1,01 | 24,47 ± 5,46 | 5,06 ± 1,94 | 8,06 ± 1,51 |
| **t<sub>max</sub> (h)** | 0,5 | 0,5 | 0,5 | 0,5 | 0,5 | 0,5 |
| **AUC<sub>0-24</sub> (µg.h/ml)** | 16,68 ± 4,41 | 9,42 ± 1,90 | 8,74 ± 2,56 | 48,40 ± 22,23 | 15,78 ± 9,10 | 17,69 ± 7,33 |
| **AUC<sub>0-∞</sub> (µg.h/ml)** | 16,87 ± 4,42 | 9,42 ± 1,90 | 8,82 ± 2,63 | 59,14 ± 24,50 | 50,90 ± 35,81 | 43,48 ± 19,92 |
| **t<sub>1/2</sub> (h)** | 4,814 ± 0,47 | 0,895 ± 0,17 | 1,871 ± 0,24 | 3,49 ± 0,69 | 8,32 ± 3,01 | 9,98 ± 5,63 |

### Effect of TW68 on Fasting-blood glucose level of *ob*/*ob* and fat induced diabetic mice

All our results showed that TW68 increases the half-life of CRY, reduces the glucose level by decreasing the transcription of *Pck1* and *G6pc* and it is well tolerated with a good oral pharmacokinetic profile *in vivo*. The genetically leptin-deficient *ob*/*ob* mouse exhibits different phenotypes, such as obesity and non-insulin dependent diabetes [50]. This animal has been used in different studies as an animal model for studying type II diabetes and related metabolic disorders [50–52]. We wanted to test if TW68 could control fasting blood glucose levels in *ob/ob* animals. Animals fasted for 4 h and 40 mg/kg/day TW68 was orally administered to male *ob/ob* mice for 5 days (n=3) along with vehicle-treated and pioglitazone control animals (n=3). Pioglitazone is used for the treatment of type 2 diabetes [53]. Their blood glucose levels were measured each day after 4 h of administering TW68 for 5 days. Animals had a reduced fasting blood glucose level compared to control animals (**Fig. 8A**). Because diabetes is caused by a high-fat diet, we want to see how TW68 affects fasting glucose levels in such a case. Animals (C57BL6/J wild type mice) were fed a diet with a 60% fat content for 3 months. The glucose levels of the animals were measured, and those with levels higher than 200 mg/dL were classified as diabetic and included in the study. These animals, similar to *ob/ob* mice, fasted for 4 h and 40 mg/kg/day TW68 was orally administered to male mice. Results indicated that the animals administered with TW68 had normal glucose levels, similar to the control animals administered with pioglitazone (**Fig. 8A**, right panel). In the same experimental setup, we showed TW68 increases the level of CRY1, which suggest enhanced its stability in liver (**Fig. 8B**). Then we isolated the total RNA and performed qPCR on liver samples. Our results indicated that the transcription levels of *Pck1*, *G6pc* and *Cry1* decreased both in *ob/ob* and wild-type mice livers (**Fig. 8C**), which is consistent with *in vitro* results. Reducing the blood glucose level. The phosphatidylinositol 3-kinase (PI3K)-AKT pathway, which mediates the majority of insulin’s metabolic effects, is activated when the insulin receptor (IR) phosphorylates insulin receptor substrate proteins (IRS proteins), which are linked to the activation of two main signaling pathways [54]. We therefore hypothesized that lowering blood glucose levels with TW68 would result in reducing insulin levels and, in turn, reducing the AKT phosphorylation. To test that total proteins from animals were subjected to Western blot analyses using anti-AKT-P. We observed a reduction of phosphorylation of AKT in animals treated with TW68 (**Fig. 8D**). We then determined the effect of fasting on glucose tolerance, C57BL/6J chow-and *ob/ob* mice were fasted for 18 (overnight) h and an oral glucose tolerance test (OGTT) was performed using 2 g/kg glucose. Blood glucose levels were monitored from the tail-tip using a hand-held glucometer in the basal state and 0, 30, 60, 90, 120 and 180 min following glucose administration. OGTT results revealed that there was no significant difference between wild type animals treated with TW68 or vehicle (**Fig. 8E**). On the other hand, the lower glucose excursion was measured in *ob/ob* mice treated with TW68 compared to *ob/ob* mice receiving vehicles (**Fig. 8E**). This suggests TW68 reduces glucose excursions in the blood.

**Figure 8:**
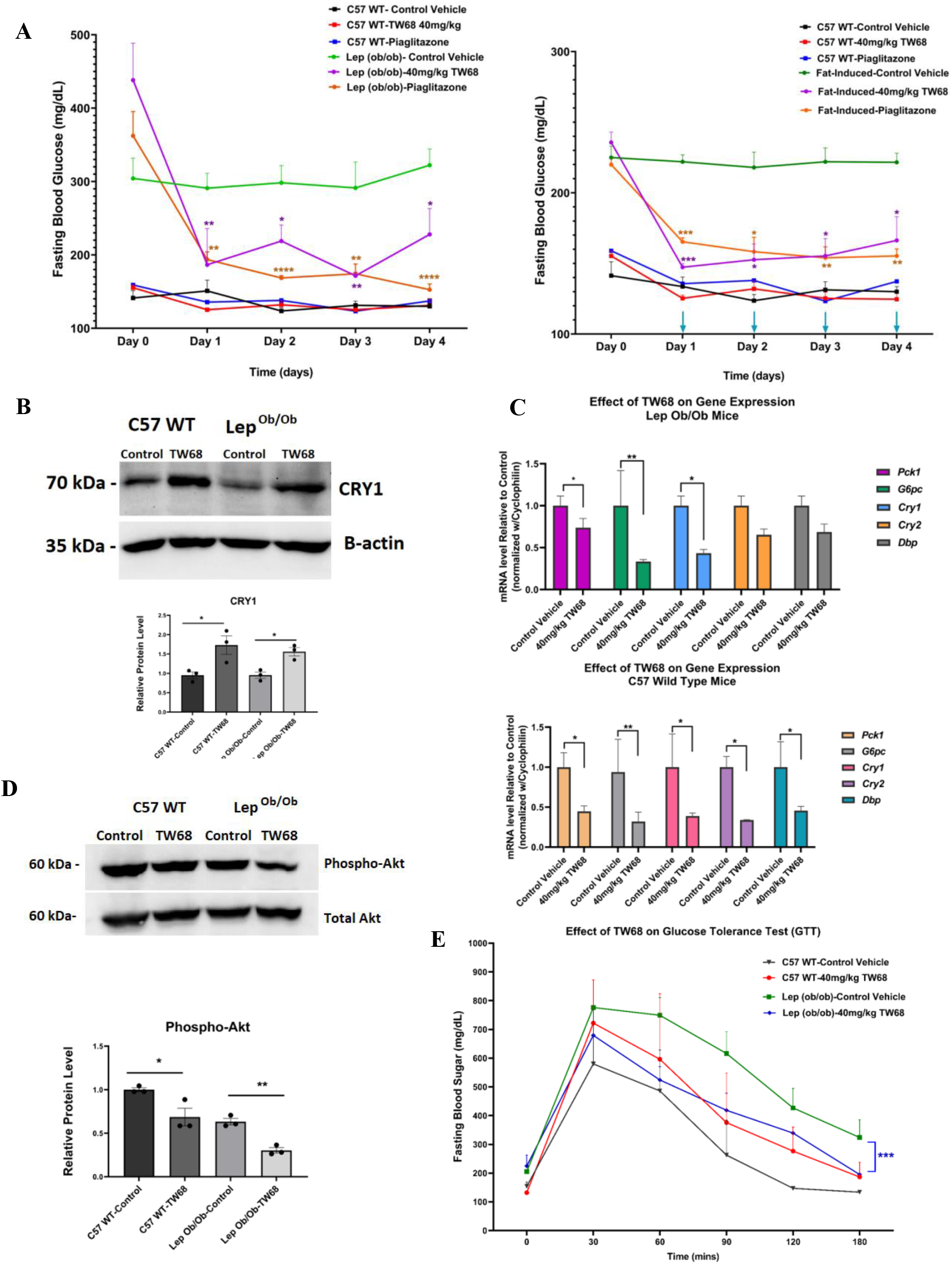
Antidiabetic Effect of TW68 on fasting blood glucose in mice. A) Left: The effect of TW68 on fasting blood glucose level of *ob/ob* diabetic animals at 40 mg/kg TW68 and 30 mg/kg pioglitazone 4 hours after oral administration in wild type and Lep (*ob/ob*) (Statistical analyses were performed by using unpaired t-test, * p<0.05, ** p<0.001, *** p<0.005, **** p<0.0001, n=3 for each group). Right: The effect of TW68 on fasting blood glucose in wild type and level fat-induced diabetic animals at 40 mg/kg TW68 and 30 mg/kg pioglitazone 4 hours after oral administration (Statistical analyses were performed by using unpaired t-test, * p<0.05, ** p<0.001, *** p<0.005, **** p<0.0001, n=3 for each group). B) Protein level of CRY1. The effect of TW68 in mice liver (WCL) 40 mg/kg TW68 or Control vehicle o.p. administered to C57BL/6J and Lep *Ob/Ob* mice. 4h after the treatment mice were sacrificed (n = 3). (Data represent the mean ± SEM, n=3, control versus TW68 treated mice by unpaired t-test *p < 0.01, **p < 0.001). C) The transcriptional level of *Pck1*, *G6pc*, *Cry1*, *Cry2*, *Dbp* in *ob/ob* and diet-induced diabetic mice. 40 mg/kg TW68 or Control vehicle o.p. administered to C57BL/6 J and Lep Ob/Ob mice. 4h after the treatment mice were sacrificed (n = 3). (Data represent the mean ± SEM, n=3, Wild type (WT) control versus TW68 treated mice by unpaired t-test *p < 0.01, **p<0.001). D) The oral glucose tolerance test (OGTT) was performed for both wild type and *ob/ob* animals. E) The effect of TW68 on oral glucose test in animals. Animals were fasted overnight and then 40mg/kg TW68 was administered. Statistical analyses were performed by using the Ordinary two-way ANOVA method by Lep (*ob/ob*) Control Vehicle versus Lep (*ob/ob*) TW68, ***p < 0.0001. Data represent the mean ± SEM, n=4 for each group)

## Discussion

The understanding of the molecular basis of circadian rhythm has been achieved through the utilization of genetic and biochemical approaches [9]. Furthermore, the relationship between the molecular circadian clock and human health has been revealed [55–58]. As a result, core circadian proteins have emerged as targets for the discovery of drug-like small molecules that can regulate their activity and, consequently, circadian oscillation [13]. Several molecules, which regulate the circadian clock through the modulation of core clock protein activity, have been identified [13, 32–34, 36, 59].

In this study, we have identified and characterized a small molecule called TW68 that stabilizes both CRYs in mammals. In a cell-based degradation assay, TW68 demonstrated a dose-dependent stabilization of CRY1-LUC and CRY2-LUC (**Fig. 1E**). When CRYs are stabilized, the increase in their repressor activity on BMAL1/CLOCK driven transcription and the period length is expected [13]. In agreement with this, longer period length was observed in TW68 treated U2OS (stably express *Bmal1-*d*Luc*) and NIH3T3 (stably express *Bmal1-* d*Luc*) cells. Our results are in agreement with previously published studies where CRY stabilizers (KL001 and KL044) have similar effects on the circadian rhythm [36, 60]. Through docking analysis of TW68 and a pull-down assay using bTW68, we found that both TW68 and KL001 compete for binding to primary pockets of CRY1.

A previous study highlighted a new role of CRY1 in which it inhibits glucagon-induced gluconeogenesis [49]. CRY1 binds to the α subunit of the G protein (Gsα) and effectively blocks GPCR-mediated increases in cAMP, thereby inhibiting gluconeogenesis. Another study using a CRY stabilizer molecule indicated that small molecules could be employed to inhibit gluconeogenesis at cellular level [49]. However, no *in vivo* studies have been reported such molecules showing their pharmacokinetics properties and efficacy in gluconeogenesis. Our *in vitro* studies provided evidence that TW68 stabilizes CRYs, resulting in reduced transcription of *PCK1* and *G6PC*, key regulators of gluconeogenesis, as well as decreased glucose levels in HepG2 cells (**Fig. 5**). Based on these findings, we hypothesized that TW68 could effectively lower blood glucose levels during fasting conditions. Consequently, our objective was to conduct pre-clinical investigations to explore the potential of TW68 for treating diabetic animals which lack control of blood glucose levels under fasting conditions. Initially, we tested toxicity and determined pharmacokinetic profile of the molecule *in vivo*. TW68 has a half-life 3.5 h in plasma and liver and well tolerated when administered orally. All these suggested that it has good pharmacokinetic profile, and good plasma and tissue distributions. In addition, TW68 is well tolerated when administered with single and repeated subacute doses. *In vivo* efficacy analysis showed that TW68 achieved to decrease fasting blood glucose level in two different conditions, genetically obese *ob/ob* mice and diet-induced obese mice, by stabilizing CRY1 and decreasing the *Pck1* and *G6pc* transcription levels. Insulin level in blood was reduced upon TW68 treatment of diabetic animals. This is most probably due to the reduction of glucose level in blood, which signals to reduce release of it from pancreas.

In conclusion, all the *in vitro* and *in vivo* results suggest that TW68 have a therapeutic potential to regulate blood glucose level by inhibiting gluconeogenesis induced by glucagon.

## Materials and Methods

### Cell Culture

HEK293T, U2OS, NIH3T3 and HepG2 cell lines were maintained in Dulbecco’s modified Eagle’s Medium (DMEM, Gibco) supplemented with 10% heat-inactivated fetal bovine serum (FBS, Gibco), 100 μg/ml penicillin, and 100 μg/ml streptomycin (Gibco). Cell cultures were grown in an 85% humidified incubator at 37°C supplied with 5% CO_2_. Cells were subcultured with Trypsin/EDTA (0.25%:0.02%, PAN Biotech) when they reach confluency.

### Cell Viability Assay

Non-toxic doses of small molecules were determined in both U2OS and HEK293T cells with MTT (3-(4,5-dimethylthiazol-2-yl)-2,5-diphenyltetrazolium bromide, AppliChem) cell viability assay [61].The cells were seeded at a density of 4×10^3^ cells per well in a 96-well plate. After 48 hours, small molecules dissolved in dimethyl sulfoxide (DMSO) were added at different doses. Following another 48 hours of incubation, the medium was vacuumed and 1 mg/mL MTT solution in DMEM was added to each well. The cells were incubated for 4 h and the formazan crystals were dissolved in 1:1 DMSO:ethanol. Using Synergy H1 Microplate Reader (Bio-Tek Instruments), the absorbance values were measured at 600nm.

### Protein Degradation Assay

We followed the protocols published in [43]. Briefly, reverse transfection of HEK293T cells was performed at a density of 4×10^4^ cells/well on a 96-well white flat bottom plates using polyethyleneimine (PEI) using pcDNA-*Cry1-Luc* and pcDNA-*Cry2-Luc*. After 24 h of incubation, small molecules (final 0.4% DMSO) were added at different doses. Then the medium was supplemented with 1mM luciferin and 10mM HEPES (pH 7.2). After 1 hour, 30 µg/ml cycloheximide was added to inhibit protein synthesis. The plate was sealed and immediately placed in Synergy H1 Microplate Reader (Bio-Tek Instruments). Every 20 minutes, the luminescence was measured for a total of 24 hours. Using GraphPad Prism 5 (GraphPad Software Inc.), the half-life was calculated from one phase exponential decay curve fit of data and dose-response curves were obtained from sigmoidal curve fitting where EC50 (half maximal effective concentration) and R^2^ (coefficient of determination) were calculated.

### Real-time monitoring of bioluminescence

The effect of the small molecules on circadian rhythm was determined using U2OS and NIH 3T3 stably expressed destabilized luciferase under the control of *Bmal1* promotor (U2OS *Bmal1-dLuc* cells and NIH 3T3 *Bmal1-*d*Luc*) [32, 34, 62]. U2OS *Bmal1-dLuc* cells were seeded in 35 mm plates at a density of 3.5×10^5^ cells and NIH 3T3 *Bmal1-*d*Luc* cells were seeded at a density of 4×10^5^ cells/plate. After 24 hours, a fresh medium containing 0.1 µM dexamethasone was added to synchronize the circadian rhythm. 2 hours later, the medium was replaced with the recording medium [DMEM (D-2902, Sigma) with 3.5 mg/ml D (+) glucose (Sigma), 0.35 mg/ml sodium bicarbonate (Sigma), 5% FBS, 10 mM HEPES (pH 7.2), 100 μg/ml penicillin, and 100 μg/ml streptomycin (Gibco)] containing freshly added 0.1 mM luciferin. Subsequently, the small molecules (final 0.4% DMSO) were added at different doses. Plates were placed in LumiCycle luminometer (Actimetrics) at 37°C and the bioluminescence was measured every 10 minutes for 7 days. Data were analyzed with LumiCycle Analysis software (Actimetrics).

### Immunoblotting

U2OS cells were seeded on 35mm dishes with a density of 3.5×10^5^ cells. After 24 hours, small molecules (final 0.4% DMSO) were added at different doses. The cells were harvested 24 hours after treatment. Cells were lysed with RIPA buffer (50 mM Tris-HCl pH 7.4, 150 mM NaCl, 0.5% NP-40, 0.5% SDS) containing 1X Protease Inhibitor Cocktail (PIC, Thermo Scientific) and 1 mM phenylmethylsulfonyl fluoride (PMSF). The samples were subjected to sodium dodecyl sulfate polyacrylamide gel electrophoresis (SDS-PAGE) and transferred to polyvinylidene difluoride membrane (PVDF Transfer Membrane, Merck-Millipore). The primary antibodies used for the detection of proteins are Anti-CRY1 (Bethyl), Anti-CRY2 (Bethyl), Anti-PER2 (Bethyl), and Anti-β-ACTIN (Cell Signaling). The secondary antibody used for CRY1, CRY2, and PER2 is an anti-rabbit-HRP antibody (Cell Signaling) and used for β-ACTIN is an anti-mouse-HRP antibody (SantaCruz). Finally, the proteins were visualized with WesternBright ECL^TM^ HRP substrate (Advansta) using ChemiDoc Imaging System (Bio-Rad).

### Real-time quantitative PCR (RT-qPCR)

Total RNA extraction was performed using NucleoSpin RNA kit (Macherey-Nagel) including DNase treatment according to the manufacturer’s protocol. The concentration and purity of RNA was measured using NanoDrop 2000 (Thermo Scientific) while the integrity was confirmed by agarose gel electrophoresis. RT-qPCR was performed as described previously [41]. The primer sequences are given in **Table 4**. mRNA levels were normalized with respect to *Rplp0* since its expression level is independent of the circadian clock mechanism. CFX Connect Real-Time PCR detection system (Bio-Rad) was used for measurement and analysis.

**Table 4:**
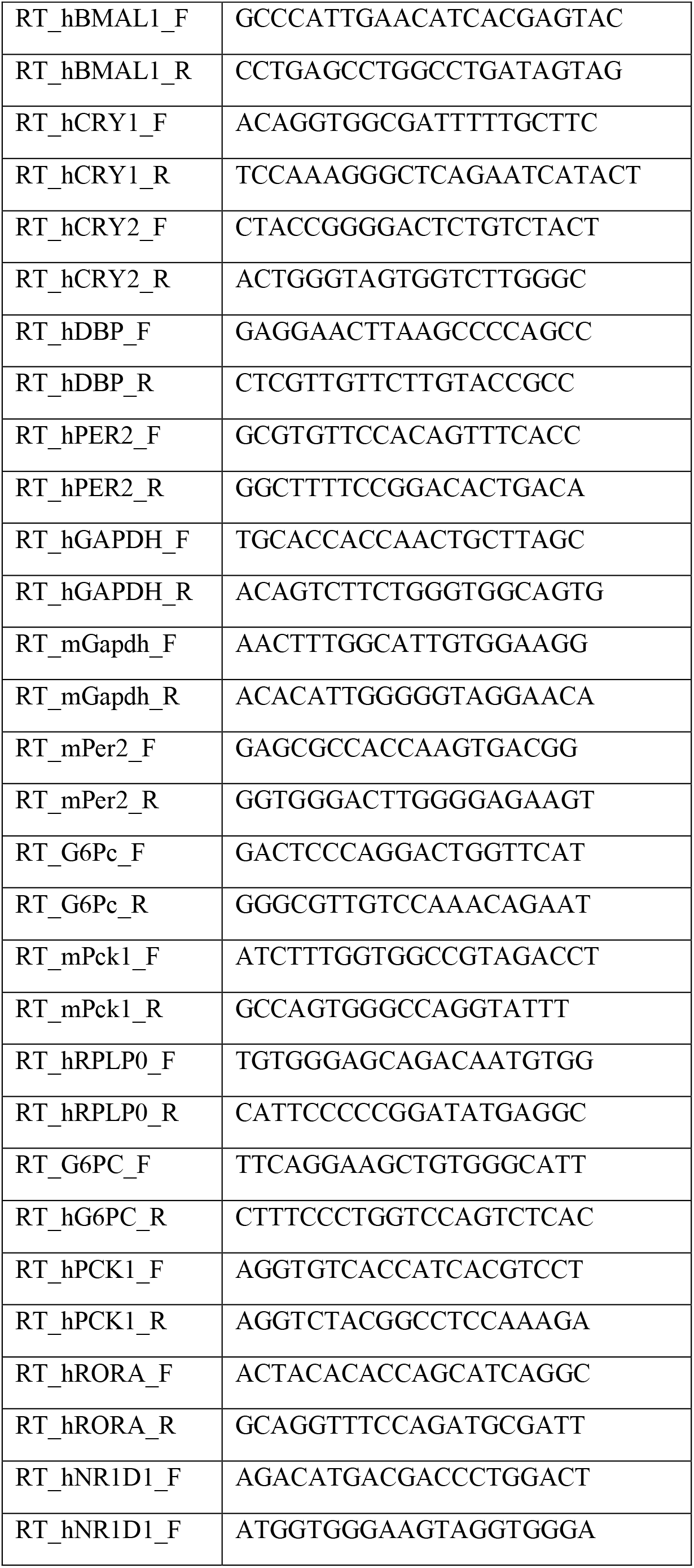
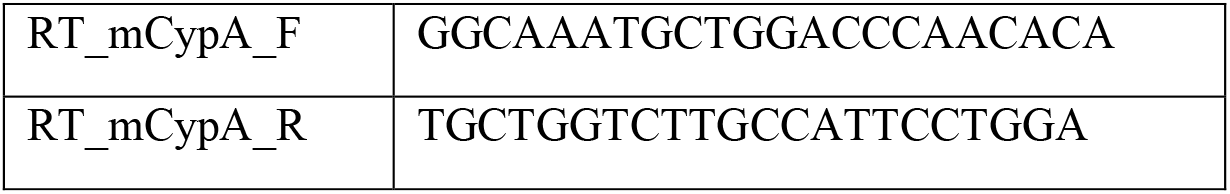
Sequence of primers for RT-qPCR.

### Real-time monitoring of *CRY* DKO U2OS bioluminescence

*CRY1*/*CRY2* double knockout U2OS were seeded on 35-mm dishes at a density of 3×10^5^ cells/plate. After 72 hours, a fresh medium containing 0.1 µM Dexamethasone was added to synchronize the circadian rhythm and after 2 hours of incubation, the medium was replaced with the recording medium containing freshly added 1 mM luciferin. Subsequently, the small molecules (final 0.4% DMSO) were added. Plates were placed in LumiCycle luminometer (Actimetrics) at 37°C and the bioluminescence was measured every 10 minutes for 7 days. Data were analyzed with LumiCycle Analysis software (Actimetrics).

### Gluconeogenesis studies

HepG2 cells were seeded on 12-well plates with a density of 2×10^5^ cells/well in low glucose (1000 mg/L) DMEM with 10% FBS, 100μg/ml streptomycin and 100μg/ml penicillin. The medium was supplemented with 2μg/ml insulin and 100nM dexamethasone. Cells were incubated for 3 hours, then the medium was changed with low glucose DMEM without FBS. Subsequently, the small molecules (final 0.4% DMSO) were added at different doses. After

18 hours, the cells were treated with 10nM glucagon. Cells used in RT-PCR analysis were harvested after 2 hours. For glucose production assay, after 3 hours, the cells were washed with warm glucose-free Krebs-Ringer bicarbonate buffer (118.5 mM NaCl, 4.74 mM KCl, 23.4 mM NaHCO_3_, 1.18 mM KH_2_PO_4_, 1.18 mM MgSO_2_, 2.5 mM CaCl_2_, and 25 mM HEPES – pH 7.6) containing 1% BSA, 100 μg/ml penicillin, 100 μg/ml streptomycin, and 0.29 mg/ml L-glutamine. Then, the cells were incubated in this buffer additionally containing 2 mM sodium pyruvate 20 mM sodium lactate for 4 hours. The glucose levels in the buffer were measured using Glucose and Sucrose Assay Kit (Sigma, MAK013).

### In vivo Studies

#### Animals and their synchronization

Male and female C57BL/6J mice, 8-12 weeks of age, weighing 18-25 g were used in the in vivo studies with TW68. Mice were obtained from Koç University Animal Research Facility (KUTTAM) and experiments were conducted in accordance with the guidelines approved for animal experimental procedures by the Koç University Animal Research Local Ethics Committee (No: 2017/08).

Mice were housed in polystyrene cages of up to four animals in a room equipped with temperature control (21 ± 2°C) and humidity (55 ± 5%). Mice were housed under 12 h of light (L) in alternation with 12 h of darkness (D) (LD 12:12) prior to any intervention and the same lighting regimen continued to the end of the experiment. Water and food were provided ad libitum throughout the experiments. TW68 or control vehicle were administered intraperitoneally and orally to mice at the same time every day (three hours after light onset).

### Preparation of TW68 formulation

TW68 was dissolved in 2.5% DMSO and 15% Cremophor EL and then diluted with 82.5% isotonic sodium chloride solution (Control Vehicle = DMSO:Cremophor EL:0.9% NaCl; 2.5:15:82.5, v/v/v) on each study day to prepare freshly, prior to intraperitoneal or oral injection. In the oral administration formulation, distilled water (dH2O) was used instead of isotonic sodium chloride solution then performed by ball-point gavage needle accordingly. Solvents were reagent grade, and all other commercially available reagents were used as received unless otherwise stated.

### Single Dose Toxicity Study: Determination of the Dose Range

The dose levels to be used in the single-dose toxicity study were selected according to the OECD Guidelines (Guidelines for the testing of chemicals, 2002). Male (n=2-3) and female (n=2-3) C57BL/6J mice were used at each dose level. Mice were treated with 50 and 300 mg/kg single doses of TW68 intraperitoneally, with one dose being used per group. Control mice (n=2 for both sexes) were only treated with the control vehicle (DMSO:Cremophor EL:0.9% NaCl; 2.5:15:82.5, v/v/v, i.p.). Careful observations of mice including body weight changes, body temperature, behavioural and clinical abnormality, and mortality were performed for 14 days, and gross necropsy of all animals was carried out at the end of the experiment. Body weight was measured every day as an index of general toxicity. TW68-induced body weight change was expressed relative to body weight on the initial treatment day. The body temperatures of the mice were recorded by the rectal homeothermic monitor (Harvard Apparatus, US) for 5 days following TW68 injection. Temperature measurements were performed at the same time each day. After the observation period, mice were exposed to isoflurane anaesthesia and blood was collected by cardiac puncture. Mice were immediately sacrificed by cervical dislocation after blood collection.

### Repeated Dose Toxicity Study: Determination of the Maximum Tolerated Dose of TW68 Upon 5-Day Administration

The single-dose toxicity data were used to assist in the selection of the doses in this repeated dose toxicity study. C57BL/6J mice (n=6 per group) were treated with 50, 75 or 150 mg/kg doses of TW68 intraperitoneally for 5 days. Control mice (n=4) were only treated with control vehicle (DMSO:Cremophor EL:0.9% NaCl; 2.5:15:82.5, v/v/v, i.p.). Throughout the study, animals were monitored for mortality, clinical signs, body weight changes, body temperature, food and water consumptions, behavior assessment and gross findings at the terminal necropsy. Body weights and body temperatures of mice were measured every day as an index of toxicity. Within 48 hours after the last dose of TW68 administration, blood was collected from mice by cardiac puncture under isoflurane anesthesia. The liver, spleen, and kidneys were removed and fixed in 10% formalin solution for histological examinations.

### Repeated Dose Toxicity Study: Determining the Subacute Toxicity of 20 mg/kg TW68 Upon 28-Day Administration

According to the results of the 5-day maximum tolerated dose determination study and histology results, TW68 dose was determined as 20 mg/kg for the next 28-day subacute toxicity study. In this study, C57BL/6J mice (n=10) were treated with 20 mg/kg dose of TW68 orally for 28 days. Control mice (n=5) were only treated with control vehicle. Following this, the previously mentioned procedure (5-day maximum tolerated dose determination study) has been conducted in the same way.

### Pharmacokinetic Studies

C57BL/6J female mice (n=4 per each time point) were treated with 75 mg/kg single dose of TW68 intraperitoneally and orally. Blood samples were collected at 0.5, 1, 2, 4, 8, 24 and 48 hours after administration of TW68 by cardiac puncture under isoflurane anesthesia. Plasma was obtained by centrifugation from heparinized tubes and stored at −80 °C until analysis. Liver, kidney, and ileum tissues were quickly removed and stored at −80 °C for further processing. For assessment of the tissue exposure of TW68, levels in these tissues were determined according to maximum concentrations of the molecule in plasma.

### Analysis of Antidiabetic Effects in Mouse Models of Type 2 Diabetes

B6.Cg-Lep^ob^/J transgenic mice from Jackson Laboratory (Strain #:000632) were used for the determination of the antidiabetic effect of TW68 in the genetic background of obesity. Female and male Lep^ob^ transgenic mice, 8-12 weeks old, weighed over 30 g were used. After a 4-h fasting, the plasma glucose levels were determined before and after oral administration of 40mg/kg TW68 molecule, 30mg/kg pioglitazone, or control vehicle via glucometer (Accu-check, Performa Nano, Roche, Switzerland) over 4 days (n=3).

### Fat-Induced Obese

The 6 weeks old C57BL6/J wild-type mice were randomly assigned to a high-fat diet feeding group for fat-induced obesity. Mice were fed ad libitum with a diet containing 60 % kcal fat (Formula #: D12492, Research Diet, USA) until the body mass increased over 30 g and plasma glucose levels above 185-190 mg/dL. For the determination of the antidiabetic effect of TW68 on fat-induced obese mice, after a 4 h fasting the plasma glucose levels were determined before and after oral administration of 40mg/kg TW68 molecule, 30mg/kg pioglitazone, or control vehicle via glucometer (Accu-check, Performa Nano, Roche, Switzerland) over 4 days (n=3).

### Oral Glucose Tolerance Test (OGTT)

Animals were undergone overnight fasting for 16 hr with access to drinking water at all times. 2 hours before the glucose loading, 40mg/kg TW68 molecule or control vehicle were administered orally. 20% (w/v) glucose solution were administered by ball-point gavage needle of 2 g glucose/kg at an oral injection volume of 10 μL/g body weight. Blood glucose level were measured from the tail vein blood sample via glucometer (Accu-check, Performa Nano, Roche, Switzerland) at different time points in a period of 3 hr (n=4).

### RNA Isolation and Quantitative Real-Time PCR (RT-qPCR) from Mice Liver

Mice were sacrificed 4 h after orally administered 40mg/kg TW68 or vehicle. QIAGEN RNeasy Mini Kit was used to isolate the mRNA. The protocol provided by the manufacturer was followed. Dounce homogenizer was used for the homogenization of the liver samples. On-column DNase treatment was performed with the DNase enzyme provided with the kit. RNA quantity was determined by Nanodrop2000 (Thermo Scientific). 500ng of each RNA sample was subjected to first strand cDNA synthesis with MMLV-reverse transcriptase (Thermo, 28025013). The mRNA expression levels were calculated by quantitative real-time PCR (RT-qPCR) with the SYBR Green. *Cyclophilin* gene was used as an internal control. All reactions were performed in biological triplicate (each triplicate with two technical replicates), and the results were expressed relative to the transcript level of *Cyclophilin* in each sample using the 2-ΔΔCT method.

### Investigation of the Protein Levels from Mice Liver by Western Blot

Mice were sacrificed 4 h after orally administered 40mg/kg TW68 or vehicle. Organs were immediately frozen in liquid nitrogen and then kept in −80⁰C for downstream analysis. Liver samples were homogenized with Dounce homogenizer with 500 μL RIPA buffer (50mM Tris, 150mM NaCl, 1% Triton-X, 0.1% SDS and Protease inhibitor cocktail, and additionally 8 mg/mL phosphatase inhibitor cocktail and 2mM Na_2_VO_4_ for phosphorylated samples) was added on 10 mg liver for the whole cell lysate analysis. Samples were incubated 10 minutes on ice then centrifuged for 10 minutes at 13000 rpm at 4⁰C. After determining and equalizing the protein amounts by using the Pierce 660 nm Protein assay (Thermo Scientific) via Biotek Synergy H1, lysates were mixed with a 4X Laemmli buffer (277.8mM Tris HCl pH: 6.8, 4.4% LDS, 44.4% (w/v) glycerol, 0.02% Bromophenol blue, 5% volume of beta-Mercaptoethanol) were boiled at 95⁰C for 10 minutes. Samples were used for SDS-PAGE and transferred to PVDF membrane (Millipore). Membrane was blocked with 5% milk solution in 0.15%TBS-Tween for 1 hour and then incubated with the following antibodies; CRY1 (Bethyl, A302-614A), Beta-Actin (Cell Signaling, 8H10D10), Phospho-Akt (Cell Signaling, 4060), Akt (Cell Signaling, 4691), HRP conjugated secondary mouse (Santa Cruz, SC-358920) or rabbit (Cell Signaling, 7074) antibodies. ECL buffer system (Advansta Western Bright) was used to visualize HRP chemi-luminescence via BioRad ChemiDoc Touch visualizer.

### Statistical Analysis

All data were expressed as means ± standard error of the means (SEM) for each studied variable. Statistical analyses were performed using GraphPad Prism for Windows (GraphPad Software, California, USA). The statistical significance of differences between groups was validated with either of these: Student’s t-test and one- or two-way analysis of variance (ANOVA), following Dunnet’s or Bonferroni post hoc tests, respectively. The type of significant test used in each experiment and their p-values were indicated in the figure legends.

## Author contributions

SS designed the study and performed the experiments and wrote the manuscript. CE performed in vitro studies, YKA and AO perform mice related experiments. AO wrote the manuscript. ZMG performed dose dependent work. TK, FA and NuriO generated CRYs knockout cell lines. OSI and MG synthesized biotinylated molecule and wrote related method section. ACG performed the analytical quantification experiments and rote related method section. NurhanO and BA performed proteomics studies. OO performed computational studies. ACT handled all mice. IHK conceived experimental design and wrote the manuscript.

## Data and materials availability

All data needed to evaluate the conclusions in the paper are present in the paper and/or the Supplementary Materials. Additional data related to this paper may be requested from the authors.

